# Investigation of mutated in colorectal cancer evolution history indicate a putative role in Th17/Treg differentiation

**DOI:** 10.1101/2022.10.06.511065

**Authors:** Norwin Kubick, Irmina Bieńkowska, Jarosław Olav Horbańczuk, Mariusz Sacharczuk, Michel Edwar Mickael

## Abstract

The MCC family of genes plays a role in colorectal cancer development through various immunological pathways, including the Th17/Treg axis. We have previously shown that MCC1 and not MCC2 play a role in Treg differentiation. Our understanding of the genetic divergence patterns and evolutionary history of the MCC family in relation to its function, in general, and the Th17/Treg axis, in particular, remains incomplete. In this investigation, we explored 12 species’ genomes to study the phylogenetic origin, structure, and functional specificity of this family. In vertebrates, both MCC1 and MCC2 homologs have been discovered, while invertebrates have a single MCC homolog. We found MCC homologs as early as Cnidarians and Trichoplax, suggesting that the MCC family first appeared 741 million years ago (Ma), whereas MCC divergence into MCC1 and MCC2 families occurred at 540 Ma. In general, we did not detect significant positive selection regulating MCC evolution. Our investigation, based on MCC1 structural similarity, suggests that they may play a role in the evolutionary changes in Tregs’ emergence towards complexity, including the ability to utilize calcium for differentiation through the use of the EFH calcium-binding domain. We also found that the motif NPSTGE was highly conserved in MCC1 but not in MCC2. The NPSTGE motif binds KEAP1 with high affinity, suggesting an Nrf2-mediated function for Nrf2. In the case of MCC2, we found that the “Modifier of rudimentary” motif is highly conserved. This motif contributes to the regulation of alternative splicing. Overall, our study sheds light on how the evolution of the MCC family is connected to its function in regulating the Th17/Treg axis.

## Introduction

Th17 play an important role in colorectal cancer. The Th17/Treg axis is a vital element of the immune system. Th17 and Treg belong to the CD4+ T cell type[1]. They share a large portion of their transcriptome [2]. However, they diverged in their functions. Th17 can be proinflammatory by producing several pro-inflammatory cytokines such as IL17A, IL17F, IL1, and IL6 [3]. Treg is an anti-inflammatory cell [4]. Treg’s primary role is to regulate adaptive immune cells through several direct and indirect methods [5]. Notably, the function of Th17 and Tregs in defending the body against disease is multi-edged. For example, over-activation of Treg can exacerbate pathogens’ colonization. In autoimmune diseases such as multiple sclerosis and other inflammatory diseases, Th17 is pathogenic, while Treg function is regulatory.

MCC family play a critical rol in Th17/Treg differentiation as well as in colrectal cancer pathology. The mutated in colorectal cancer (MCC) gene was identified based on its linkage to the susceptibility locus for familial adenomatous polyposis on chromosome 5q. familial adenomatous polyposis is a rare hereditary cancer predisposition disease presented by hundreds to thousands of precancerous colorectal polyps (adenomatous polyps), that, if left untreated develop into colorectal cancer. Since then MCC mutations have been identified in various cancers including colon cancer, eosesphgus cancers and breast cancers[6]. Recently it was shown that MCC gene could be contributing to cancer progression through dysregulating the WNT pathway[7]. The role of Th17/Treg axis dysregulation in colorectal cancer is widely documented. It has been shown that Th17 through the production of IL17,ILF case increase colorectal progression, in contrary to Treg role. It has been demonstrated that Treg suppression of Th17 in colorectal cancer can reduce cancer progression and improve prognosis [2]. We have recently found that MCC1 but not MCC2 contributes to the differentiation of naïve T cells into Tregs.

The role of evolution in directing MCC family in manipulating Th17/Treg in colorectal cancer is unknown. The MCC family plays a variety of molecular roles. Recent research has revealed that the nuclear factor-B (NF-B) pathway and cell cycle regulation, two crucial cellular processes related to carcinogenesis, may be affected by the wild-type MCC protein (Matsumine et al., 1996). The activation and translocation of NF-B to the nucleus is part of a signal-transduction pathway that forms the basis for several physiologic and pathologic processes, including cancer. Additionally, it was reported that MCC1 negatively regulates cell cycle progression. The orthologous gene in the mouse expresses a phosphoprotein associated with the plasma membrane and membrane organelles, and overexpression of the mouse protein inhibits entry into the S phase. Another protein known as USHBP1 (USH1 Protein Network Component Harmonin Binding Protein 1) was reported to belong to the MCC family. However the phylogenetic relationship between the two proteins is currently unknown.

In this report, we analyzed the evolutionary history of the MCC family using a host of phylogenetic tools. Previously we found, upon inspection of microarray/RNAseq data that compares Th17 and Treg differentiation, that MCC1 and not MCC2 is upregulated in Treg but not in Th17 along with a host of genes that are associated with cell cycle regulation. We found that this family could be grouped into two subfamilies, namely; MCC1 and MCC2. These two families diverged from a single MCC homolog during the Cambrian explosion (e.g., during Vertebrate emergence), when two rounds of genome duplication occurred. However, the invertebrates MCC first appeared in Trichoplax. Our investigation indicates that the likely parent family for the MCC family is a protein family containing an EF-hand domain. We investigated the conserved domains of the two families. The main building blocks of MCC in vertebrates are two domains of MCC-PDZ. This structure is found in Cnidaria and partially conserved in other invertebrate species indicating functional conservation between invertebrates and vertebrates. Moreover, motif inspection suggests thatMCC1 and MCC2 families could be playing a primary role in cell cycle regulation. However, we found that the motif NPSTGE which is known to play a role in interaction with KEAP1, only exists in MCC1. Our finding suggest that MCC1 could be enhancing Treg differentiation through inhibiting KEAP1 effect.

## Methods

### 2.1 Database search

The aim of this study was to examine the relationship between the molecular evolution of the MCC family and its functions. We reasoned that due to the diversity of the two protein subfamilies, studying protein sequences rather than DNA sequences might be more informative (MCC1 and MCC2). Furthermore, we selected 12 species and genera that span more than 500 million years to ensure that our analysis is a fair representative of the evolutionary history of the MCC family. BLASTP searches were conducted using human MCC1 and MCC2 protein sequences against the proteomes of Chimpanzee (Pan troglodytes), House mouse (Mus musculus), common wombat (Vombatus ursinus), platypus (Ornithorhynchus anatinus), red junglefowl (Gallus gallus), Zebrafish (Danio rerio), sea squirt (Ciona intestinalis), Arthropoda, Spiralia, and cnidaria, Trichoplax and sponge (Amphimedon queenlandica)[8]. Whenever one protein possessed more than one transcript, only the longest transcript was used in the analysis[9]. Sequences were selected as candidate proteins if their E values were<;1e-10[10]. Conserved domains were investigated using the CDD function on the NIH website (Accessed on 25 August 2022), PFAM and HMRR[11]. Based on the consensus domains agreed upon by the these three domain-predicting methods, sequences were filtered based on possessing conserved domains that are homologous to human MCC1 and MCC2 domains, respectively.

### 2.2 Alignment and phylogenetic analysis

The phylogenetic investigation was performed in three stages. First, MCC family amino acid sequences were aligned in Seaview using MUSCLE [12]. After that, we employed PhyML to determine the best phylogenetic tree to represent the inter-relationships among the MCC family homologs. We used the LG model with empirical values calculated for amino acids equilibrium frequencies [13][14]. Invariable sites and across-site rate variation were calculated using an optimized algorithm.

### 2.4 Functional divergence estimation and motif search

We used Sequence Harmony and Multi-RELIEF to identify residues that could be correlated with functional differences within the MCC subfamilies that caused the difference in function in the Th17/Treg axis. Sequence Harmony compared the two groups of sequences (e.g., MCC1 and MCC2 families) to identify the variable amino acids and their distribution frequency. Positions where the amino acid compositions of the two groups differ possess low score values. Hence, a score of zero indicates that the amino acids at a given position are different between the two investigated groups. A score of one indicates that the amino acid compositions are nearly identical. Multi-RELIEF predicts the residue specificity of residues, by doing two comparisons, the first is between each sequence and its nearest homolog within the same group. The second comparison is done between each sequence and its nearest homolog in the second group. A reside is deemed specific if it has a high score of specificity in at least of the of two comparisons. We conducted an extensive motifs search using ELM server http://elm.eu.org/ with a motif cut-off value of 100. We also conducted motif search using www.genome.jp[15].

### 2.5 Positive selection analysis

To investigate the selection process that governed the MCC family evolution, we utilized the maximum likelihood approach method. We back-translated the downloaded protein sequences using the EMBOSS server (https://www.ebi.ac.uk/Tools/st/emboss_backtranseq/ (accessed on 5 September 2022) to estimate the cDNA of the investigated sequences[16]. After that, we used CODEML-PAML (V4.4) to estimate the substitution rate ratio (ω) given by the ratio between nonsynonymous (dN) to synonymous (dS) mutations. We utilized four models, namely; general (basic), branch, branch-site and sites. The main difference between these for models is their level of investigation. While the global models assumes a constant ω ratio for all the tree investigated, the branch model calculates two ω values (the branch investigated and the tree, respectively). In case of the branch-site model calculates ω values for each nucleotide on a specific branch, while sites models estimate ω values for each nucleotide in the alignment. Statistical significance is calculated based on χ2 test given by the equation P value □ = □χ2 (2*Δ(ln(LRTmodel) □ – □ ln(LRTneutral)), number of degrees of freedom) (ref)[8][17].

## Results

### 3.1 Phylogenetic analysis

#### Phylogenetic analysis indicates that two rounds of duplications resulted in the divergence of MCC1 and MCC2

**W**e downloaded human MCC1 and MCC2 protein sequences from the GEO protein repository. We utilized the Blastp server to acquire homologous proteins for MCC1 and MCC2 among twelve species (Supplementary file 1 includes fata files used). We employed Seaview to perform the multiple sequence alignment using the Muscle algorithm (figure 1a,1b, and 1c). After that, we constructed the phylogenetic tree using the PyML method utilizing the LG model. Our results indicate that only one homolog of the MCC family is found in invertebrates. Our research identified a putative MCC homolog (e.g., XP_019849192.1) Amphimedon queenslandica. However, Blastp E-value was lower than our threshold of 1E-10. Thus that sequence was not accepted. Two homologs were found in all vertebrate species investigated. The Bony fish seem to be the first vertebrate to possess a pair of MCC homologs. These results indicate that the divergence of the MCC1 and MCC2 homologs occurred at the 2R stage of the Cumbrian explosion. We found that, MCC1 in vertebrates contains two MCC-PDZ domains. Cnidaria, Spiralia and Arthropoda have a similar structure composition, Tunicate and Trichoplax contain only an MCC-PDZ domain. MCC2 platypus, Vombatus, mouse, chimp and humans have two MCC sequences. Birds and bony fish contain 1 MCC-PDZ domain. Interestingly, the results of domain prediction for both MCC1 and MCC2 differed between the three servers used. Mainly NIH server predicted the existence of unique SMC domains in various MCC1 and MCC2 sequences investigated (Supplementary file 2 included NIH server results). Interestingly both PFAM and HMM as well as other domain prediction servers did not identify this unique domain in any of the species investigated. According to NIH server this domain is homologues to at least parts of the SMC gene families. We investigated further this line by BlastingP MCC1 sequences against respect species identified using the NIH server. However Blastp results did not indicate any homology between the two groups (MCC sequences and the SMC families). Thus we did not take into consideration the SMC domain in our downstream analysis. However experimental validation could shed light on this controversial aspect.

**Figure 1.**
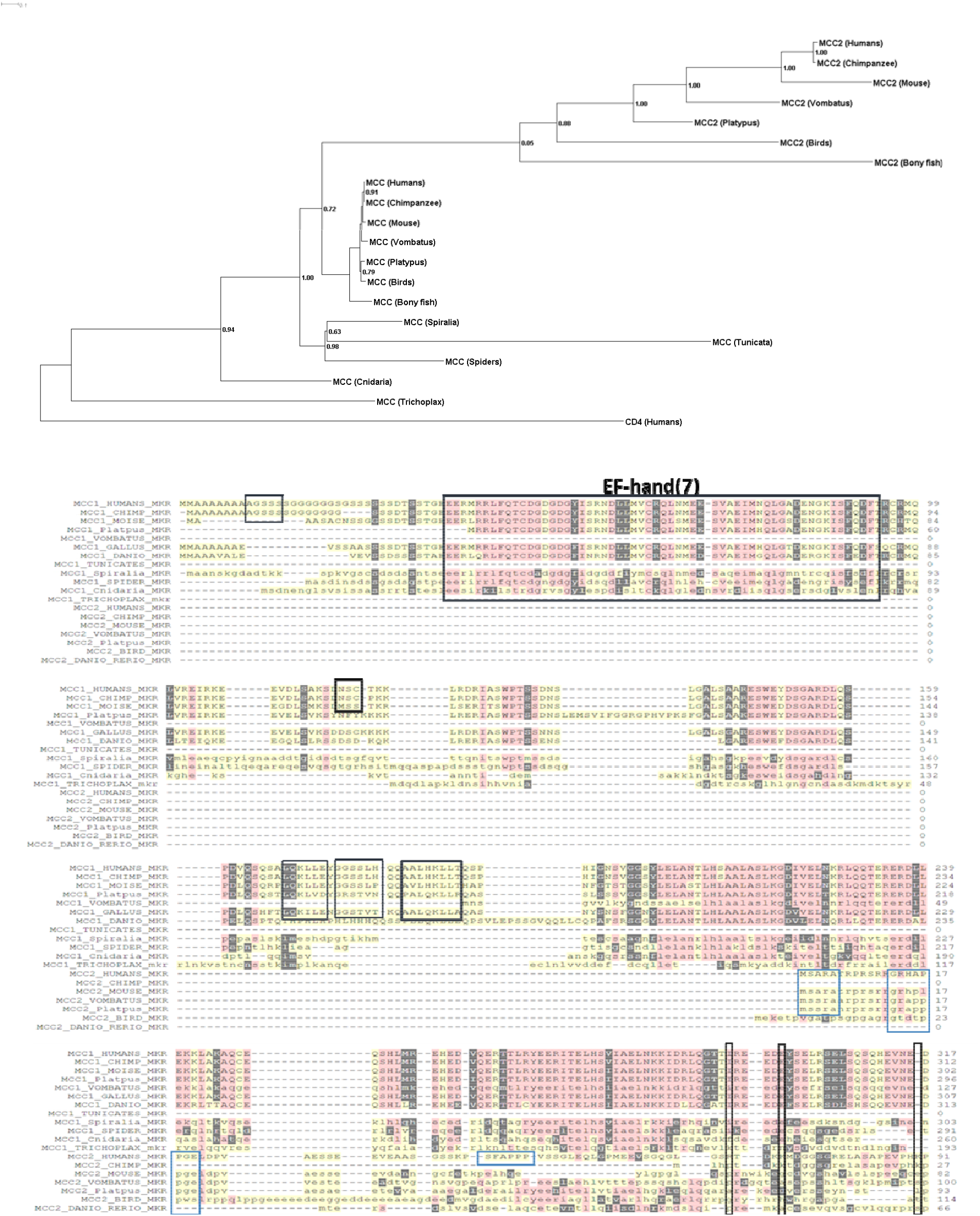
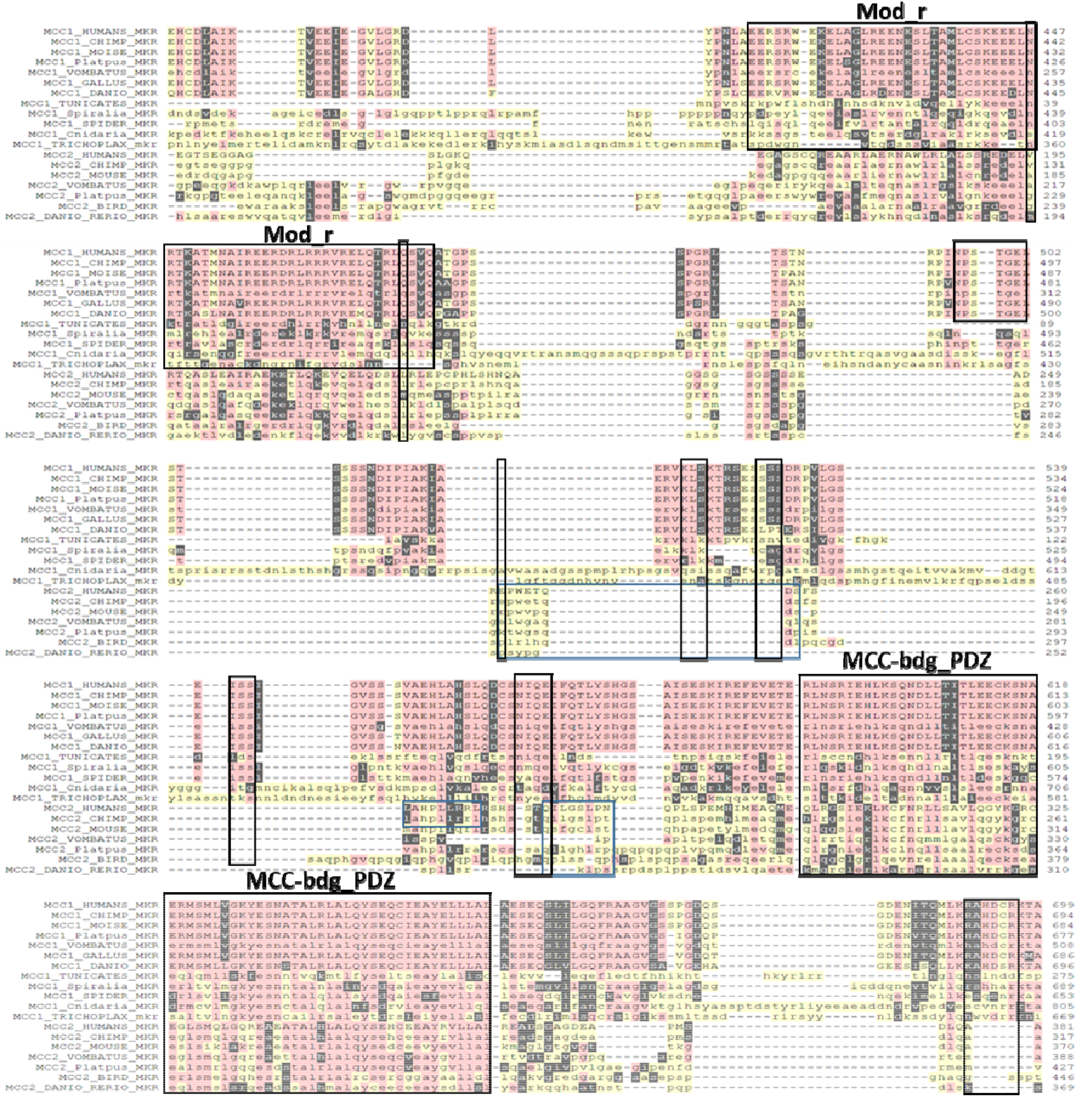
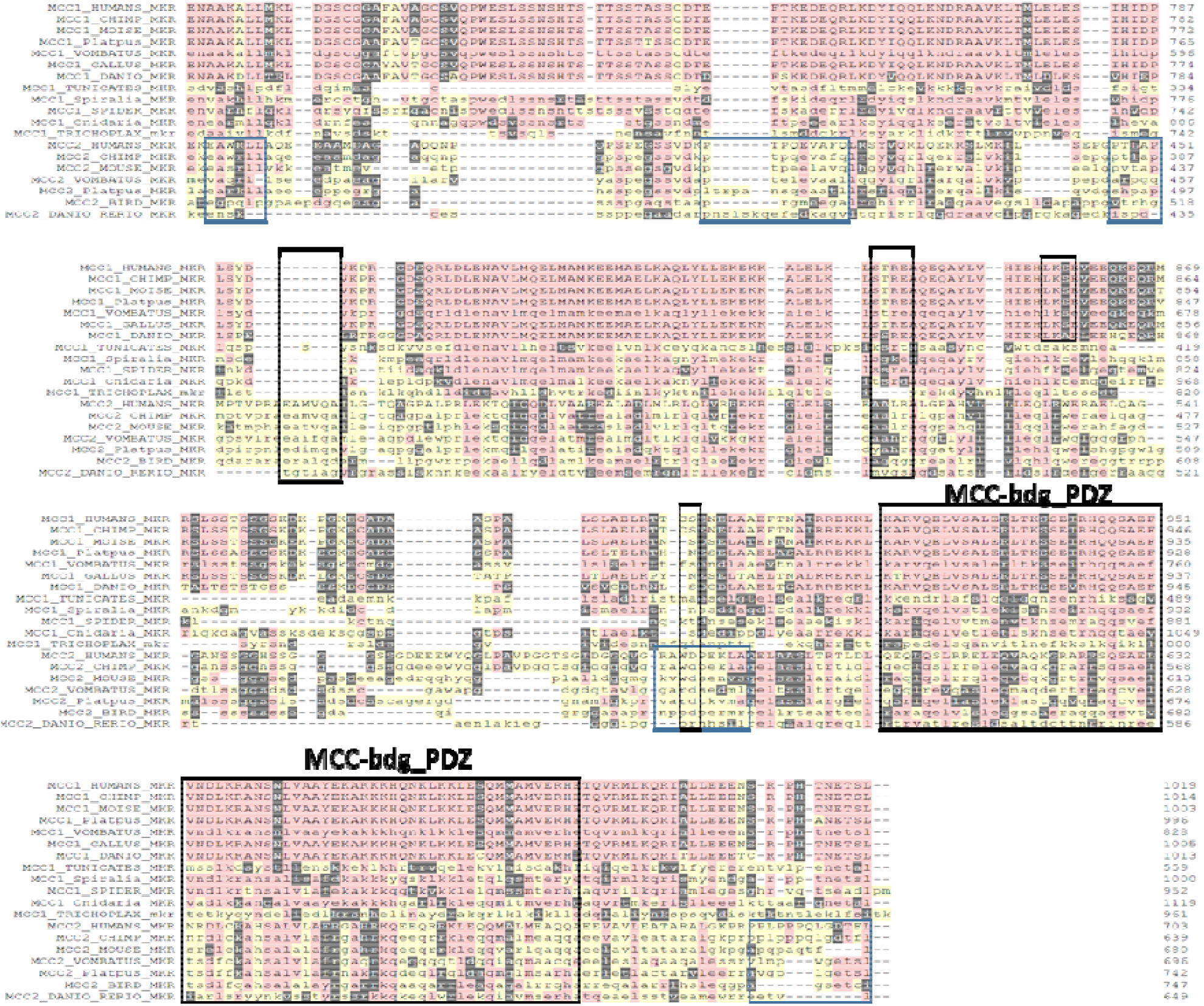
Phylogenetic analysis of the MCC family. (1A) Multiple sequence alignment of the MCC family members showing a high degree of homolog with each family. (1B) The two rounds of duplication are clear within the family species. Only one homolog for the MCC family appears in vertebrates. During the Cumbrian explosion of vertebrates, gene duplication occurred giving rise to two distinct homologs MCC1 and MCC2.

### 3.2 Ancestral sequence reconstruction and network split results

Our Ancestral sequence reconstruction indicates that the MCC closest-most ancient homolog is an EH hand domain-containing protein. We used Mega-x ancestral reconstruction function with the default setting to reconstruct the ancestral sequence of the MCC family (i.e, MCC1 and MCC2 homologs). The sequence reconstruction was based on our generated phylogenetic tree (Figures 1b and 2a). We searched for homologs for the generated ancestral sequence using BLASTP and HMM search server. The highest scoring results were EF-hand domain-containing protein, UBZ1 type domain-containing protein, ETS domain-containing protein, and ABHD8. The results and the reconstructed sequence were fed into the SplitsTree program. The results show that the nearest homolog to the ancestral sequence is EF-hand domain-containing protein (figure 2b).

**Figure 2.**
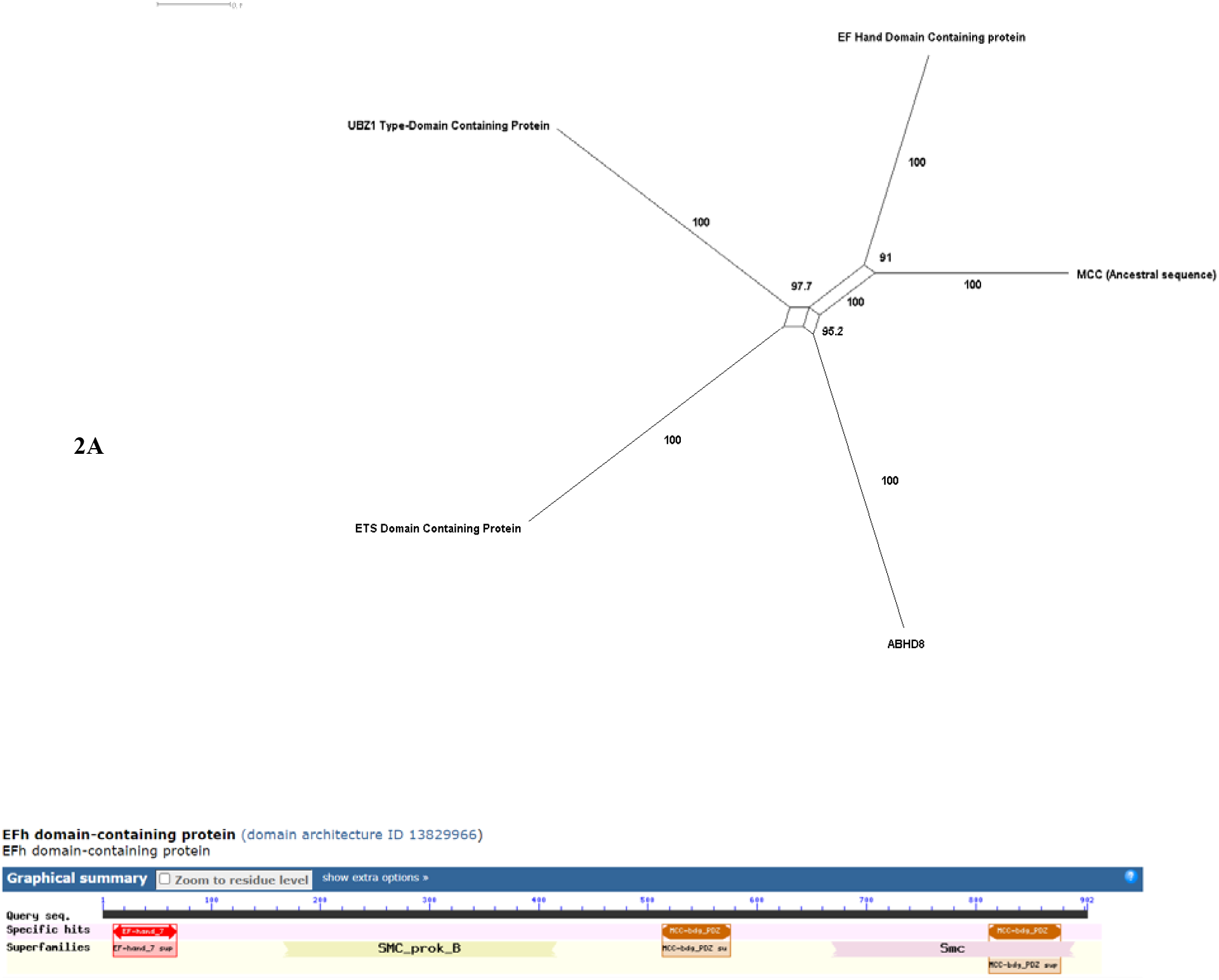
Ancestral sequence reconstruction of the MMC family. (2A) The nearest homolog for the reconstructed ancestral sequence resembles that of an EF-hand domain-containing protein. (2B) THE EF-hand domain-containing protein structure contains two MCC-PDZ domains and an SMC domain.

#### 3.4.1 Sites of Functional divergence

A study of site-specific shifts in evolutionary rates following gene duplication was performed using sequence harmony. Multiple sites seem to play a primary role in the specificity of divergence between MCC1 and MCC2. One of the important residues that seem to play a critical role in determining functional specificity between the two groups is 949-D. All the MCC1 group has a D residue, while MCC2 has R, H, or K. this residue lies in the second SMC domain of the MCC1 group and the sole SMC domain in the MCC2 group. Also in the same region, there is another critical residue (952-N) given by N/T for MCC1 while MCC2 possess K.

#### 3.4.2 Motifs

**we investigated differnetial motifs between MCC1 and MCC2 squence (Table 2)**.

**Table 1.** Positive selection Models investigated.

| Model | w |
| --- | --- |
| general | 0.45728, P-Value is < .00001 |
| branch | w value was 0.89 p-value>0.005 |
| Branch- site | w value was 0.64046 p-value>0.005 |
| sites | No sites with w>1 |

**Table 2.** Functional divergent sites.

| Pos | Entropy |  |  |  | SH | Rnk | Consensus |  |
| --- | --- | --- | --- | --- | --- | --- | --- | --- |
| Ali | A | B | AB | rel. |  |  | A | B |
| 951 | 1.21 | 0.00 | 1.71 | 1.05 | 0.00 | 11 | Adlv | - |
| 949 | 0.82 | 1.66 | 2.08 | 1.05 | 0.00 | 11 | Rkl | -ags |
| 948 | 0.92 | 2.13 | 2.31 | 1.05 | 0.00 | 11 | Fc | -gnqr |
| 947 | 1.19 | 1.84 | 2.38 | 1.05 | 0.00 | 11 | Qne | ATlp |
| 945 | 1.21 | 1.84 | 2.39 | 1.05 | 0.00 | 11 | Lefr | DPav |
| 956 | 1.95 | 1.66 | 2.79 | 1.05 | 0.00 | 11 | S-gil | Pnrt |
| 1091 | 0.00 | 1.15 | 1.37 | 1.05 | 0.00 | 9 | - | Epv |
| 1093 | 0.00 | 1.15 | 1.37 | 1.05 | 0.00 | 9 | - | Gdk |
| 1092 | 0.00 | 1.38 | 1.46 | 1.05 | 0.00 | 9 | - | Pdl |
| 1089 | 0.41 | 1.15 | 1.63 | 1.05 | 0.00 | 9 | Ed | -ap |
| 1088 | 0.41 | 1.15 | 1.63 | 1.05 | 0.00 | 9 | Lv | -ak |
| 1084 | 0.82 | 1.56 | 2.04 | 1.05 | 0.00 | 9 | Tav | PLS |
| 1085 | 1.55 | 1.15 | 2.35 | 1.05 | 0.00 | 9 | Mvip | -lq |
| 1106 | 0.00 | 0.59 | 1.17 | 1.05 | 0.00 | 8 | - | Et |
| 1110 | 0.00 | 0.59 | 1.17 | 1.05 | 0.00 | 8 | - | Qa |
| 1111 | 0.00 | 1.15 | 1.37 | 1.05 | 0.00 | 8 | - | Adg |
| 1107 | 0.00 | 1.66 | 1.56 | 1.05 | 0.00 | 8 | - | Adgr |
| 1109 | 0.00 | 1.84 | 1.63 | 1.05 | 0.00 | 8 | - | VLfm |
| 1108 | 0.41 | 1.95 | 1.93 | 1.05 | 0.00 | 8 | -s | IMTa |
| 1114 | 1.73 | 1.38 | 2.55 | 1.05 | 0.00 | 8 | P-nt | Gem |
| 778 | 0.41 | 0.59 | 1.43 | 1.05 | 0.00 | 7 | Sk | -d |

**Table 3.**
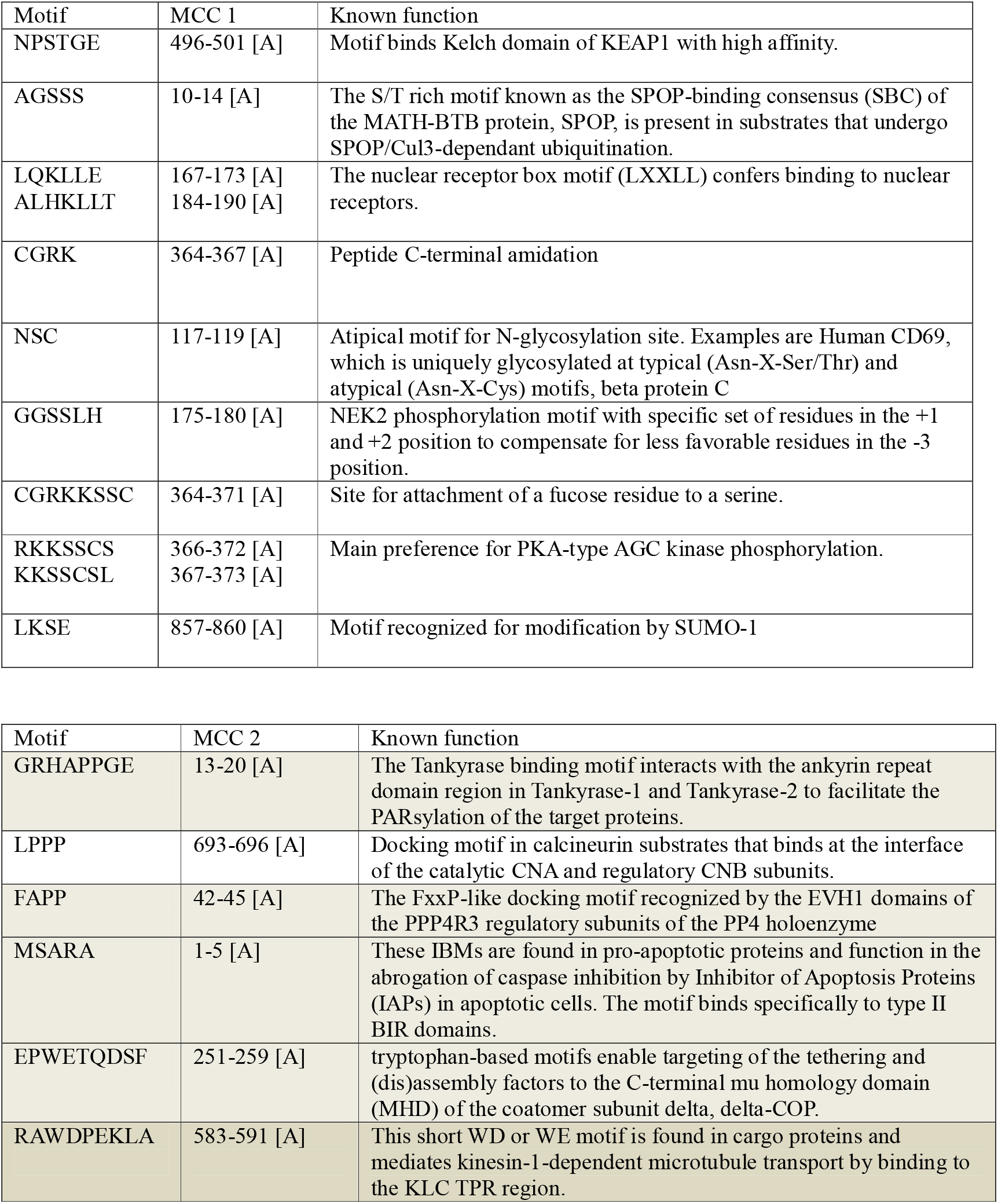
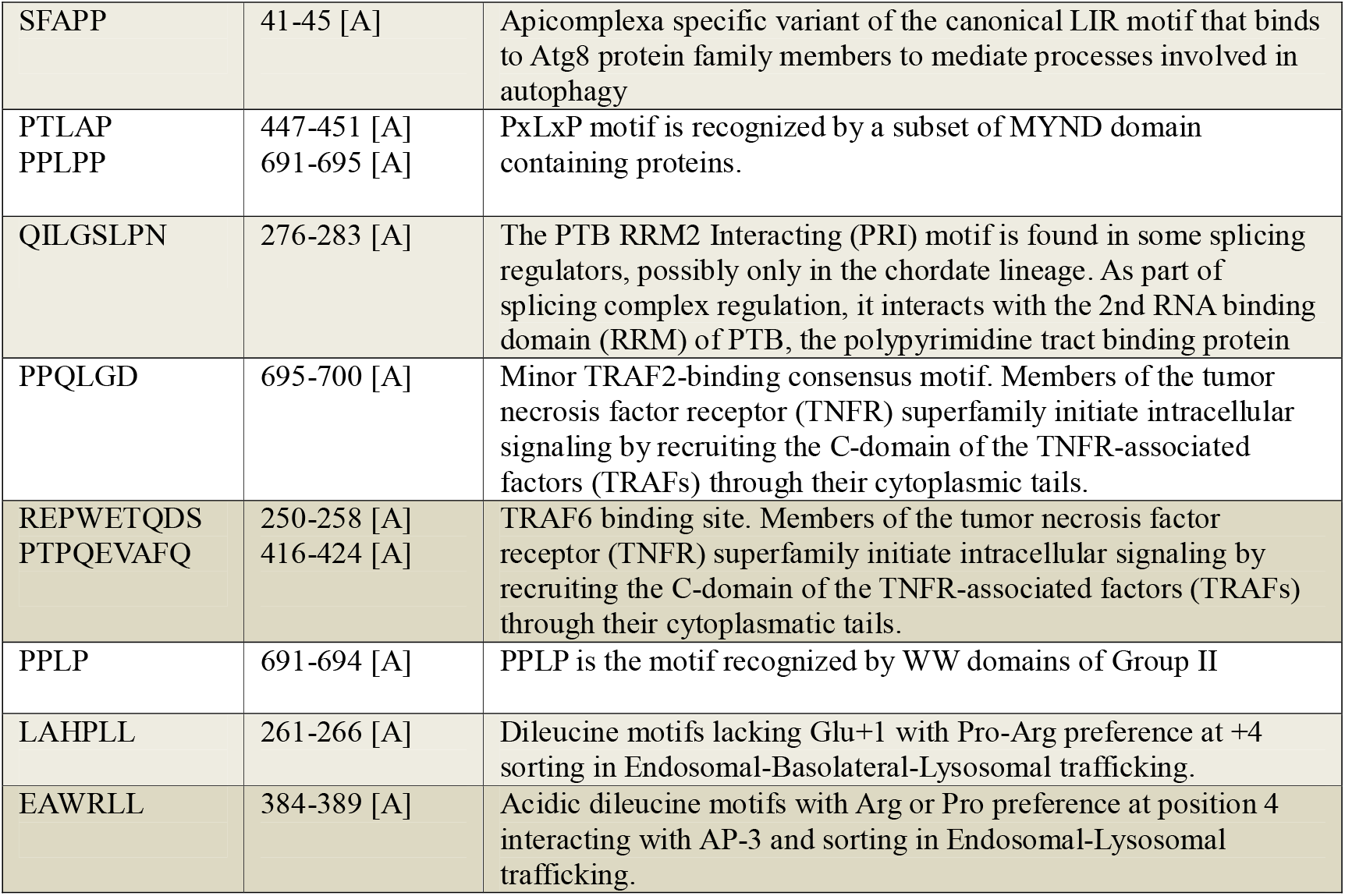
Motifs analysis and their location.

Several importnat motifs that exist only in one of teh groups sich as ERRSLMKIL that exists in primates MCC2 and not MCC1 (figure 1). Another motif is GRHAPPGE and it functions as The Tankyrase binding motif interacts with the ankyrin repeat domain region in Tankyrase-1 and Tankyrase-2 to facilitate the PARsylation of the target proteins. It mainly plays a spindle pole during cell cycle. Notably, we found in MCC1 but not in MCC2 the motif NPSTGE. this motif is binds to the Kelch domain of KEAP1 with high affinity. This high affinity motif is required for the efficient recruitment of target proteins to the Cul3-based E3 ligase. KEAP1 regulate the function of NRF2.

### 3.5 Positive selection

We conducted a positive selection analysis for the MCC family. Our results indicate that the MCC family did not seem to have evolved under significant positive selection. Conversely on several levels of analysis namely, general, branch and sites, our result indicate that MCC evolutionary pattern followed a negative selection. (i) for the global model we used M0 to denote the possibility of positive selection over the whole tree. W value calculated using PAML is 0.45728, P-Value is < .00001 (ii) Similarly, For the branch values, for MCC1 vertebrates, w value was 0.89 suggesting purifying selection, how it was not statistically significant. Also, In the case of the MCC2 branch, w value was 0.64046 without beings statistically significant. On the site level, we did not detect any sites that were subjected to significant positive selection (i.e, W>1).

## 4. Disscusion

### 4. 1 Phylogenetic analysis and origin of the MCC family

Our results indicate that the MCC family is of ancient origin. We located MCC1 and MCC2 homologs in vertebrates (figure 1). However, we found various MCC homologs in invertebrates, including Trichoplax and cnidarians. Positive section analysis did not reveal sites that were subjected to significant positive selection, indicating overall conservation of structure between the two families and within each family. We propose that MCC is linked to the EFh binding domain families. The structure of the MCC1 in vertebrates includes two MCC-PDZ domains, Cnidaria possess similar domains (figure 1). Conversely, other invertebrates possess different combinations of these domains. Our results indicate that the ancestral sequence of the MCC family is highly similar to that of the EFh binding domain (figure 2). The EFh binding domain was shown to bind calcium ions. EF-hand domain binding to calcium ions is involved in various functions, such as buffering calcium in the cytosol, signal transduction, and contraction of fibers [18][19][20].

#### Motifs and SH sites predict evolution of function between MCC1 and MCC2 family

Our results indicate a divergence of function between MCC1 and MCC2. We have found nine motifs conserved in MCC1 but not in the MCC2 family. Additionally, 14 motifs were specific only to MCC2. However, it could be noted that in the case of MCC1, these functional motifs were highly conserved in most of the species investigated. For example, the Mod(r)) represents a conserved region approximately 150 residues long within several eukaryotic proteins that show homology with the Drosophila melanogaster Modifier of rudimentary (Mod(r)) proteins. The N-terminal half of Mod(r) proteins are acidic, whereas the C-terminal half is basic[21]. This motif is highly conserved within all MCC1-expressing species. Interestingly, one site given by “N-447” residue is highly functional specific. As a result, the Mod (r) proteins may play a primary role in the function specification between MCC1 and MCC2. Another motif that seems to be contributing to the functional specificity of MCC1 is CGRKKSSC (364–371). This motif is known to be a site for the attachment of fucose residue to serine. This motif has been implicated in Notch signaling, where it was shown that the O-fucose moieties can act as a substrate for Fringe. Fringe is a known modifier of the Notch function [22][23]. However, the mechanism of interaction between MCC1 and notch signaling is still unknown. In the case of MCC2, the motif QILGSLPN is highly conserved. QILGSLPN was shown to play a role in alternative splicing. It was demonstrated that the PTB RRM2 Interacting (PRI) motif interacts with the 2nd RNA binding domain (RRM) of PTB (polypyrimidine tract binding protein). PTB acts to regulate mRNA splicing, polyadenylation, 3′ end formation, internal ribosomal entry site-mediated translation, localization, and stability. This indicates that MCC2 could be playing a role in regulating the alternative splicing of RNA [24][25].

### 4.2 A putative role for MCC1 but not MCC2 in Treg differentiation

Our research suggests that MCC1 and not MCC2 could be contributing to Treg function. In our recent publication, using microarray analysis of the Th17/Treg pathway, we revealed that several genes that regulate the cell cycle are upregulated in Treg but not Th17. MCC1 was upregulated in Tregs but not in Th17, while MCC2 was not deferentially expressed between Th17 and Tregs. Several other genes linked to controlling the cell cycle were also deferentially expressed. For example, we found that WWP2, which is known to control the cell cycle, is upregulated in Treg but not in Th17. Additionally, IL6, as expected, was not regulated in Tregs. Interestingly, wwp2-/-cells show higher IL6 levels[26]. Importantly, our results indicate that MCC1 and not MCC2 possess the motif **NPSTGE**. This motif binds to the Kelch domain of KEAP1 with high affinity. KEAP1 is one of the main regulators of NRF2. NRF2 is a known antioxidant gene that plays a role in cell cycle regulation. Additionally, we could hypothesize that MCC1 could be activating the Kelch domain of KEAP1 independently of Nrf2 [27][28]. Interestingly, Tregs are known to utilize calcium to inhibit Th17. Tregs’ mediated regulation of conventional T cells is the suppression of calcium signaling. It has been proposed that calcium inhibition of calcineurin inhibits NfkB and NFKb via an IKK-mediated pathway. Based on the existence of the EFh-binding motif in MCC1 and not in MCC2, and the upregulated expression of MCC1 and MCC2, our results hint at a putative role for MCC1 in Treg calcium production, in particular through inhibiting calcineurin-mediated pathways[29][30]. Thus, taken together, our results indicate that MCC1 is a member of a large network of genes that are capable of specifically controlling the Treg cell cycle but not Th17.

## Acknowledgments

We would like to acknowledge Macarious Abraham and Meriam Yoakim for their continuous support.

## Notes

### Competing Interest Statement

The authors have declared no competing interest.

