## Supplementary material 1 for "Investigation of mutated in colorectal cancer evolution history indicate a putative role in Th17/Treg differentiation"

### Sequence Harmony

Generated on Wed 24 Aug 2022 12:14 by SeqHarm version 1.1
from the Sequence Harmony Webserver at http://www.ibi.vu.nl/programs/seqharmwww/
Please cite:

K. Anton Feenstra, Walter Pirovano, Klaas Krab and Jaap Heringa
*Sequence Harmony: Detecting Functional Specificity from Alignments*
Nucl. Acids Res., 2007, web server issue, accepted

and

Walter Pirovano*, K. Anton Feenstra* and Jaap Heringa
"*Sequence Comparison by Sequence Harmony Identifies Subtype Specific Functional Sites*"
Nucl. Acids Res., 2006, Vol. 34, No. 22 6540-6548
* joint first authors.

Group A: 12 sequences, length 1164
Sequences: MCC_HUAMNS_MKR, MCC_CHIMP_MKR, Vombatus_MKR, Platypus_MKR, MCC_MICE_MKR, MCC_GALLUS_MKR, MCC_DANIO_RERIO_MKR, Tunicata_MCC_MKR, MCC_SP
Group B: 7 sequences, length 1164
Sequences: MCC2_HUMANS_MKR, MCC2_CHIMP_MKR, MCC2_MOUSE_MKR, MCC2_VOMBATUS_MKR, MCC2_Platpus_MKR, MCC2_BIRD_MKR, MCC2_DANIO_RERIO_MKR
Cutoff: 0.15.

Archive of all output files

#### Results for your input

Raw Table

Selected 175 positions below cutoff (0.15)
**SH:** 0.00 0.03 0.06 0.09 0.12 0.15 0.32 0.49 0.66 0.83 1.00

| Pos | Entropy | | | | SH | Rnk | Consensus | |
|---|---|---|---|---|---|---|---|---|
| Ali | A | B | AB | rel. | | | A | B |
| 547 | 0.41 | 0.00 | 1.21 | 1.05 | 0.00 | 9 | Ai | - |
| 552 | 0.41 | 0.00 | 1.21 | 1.05 | 0.00 | 9 | Lp | - |
| 551 | 1.21 | 0.00 | 1.71 | 1.05 | 0.00 | 9 | Krsv | - |
| 548 | 1.21 | 0.00 | 1.71 | 1.05 | 0.00 | 9 | Edkn | - |
| 554 | 1.21 | 0.00 | 1.71 | 1.05 | 0.00 | 9 | Kprt | - |
| 550 | 1.21 | 0.00 | 1.71 | 1.05 | 0.00 | 9 | Vkms | - |
| 549 | 1.21 | 0.00 | 1.71 | 1.05 | 0.00 | 9 | Regl | - |
| 553 | 1.28 | 0.00 | 1.76 | 1.05 | 0.00 | 9 | SKm | - |
| 555 | 1.58 | 0.00 | 1.95 | 1.05 | 0.00 | 9 | Tcflm | - |
| 608 | 0.41 | 0.00 | 1.21 | 1.05 | 0.00 | 7 | Sg | - |
| 610 | 1.21 | 0.00 | 1.71 | 1.05 | 0.00 | 7 | Gekn | - |
| 605 | 1.04 | 0.59 | 1.82 | 1.05 | 0.00 | 7 | Edg | Qk |
| 609 | 1.42 | 0.00 | 1.85 | 1.05 | 0.00 | 7 | Ilem | - |
| 611 | 1.55 | 0.00 | 1.93 | 1.05 | 0.00 | 7 | Vlce | - |
| 607 | 1.58 | 0.00 | 1.95 | 1.05 | 0.00 | 7 | Sadet | - |
| 304 | 1.58 | 1.84 | 2.63 | 1.05 | 0.00 | 6 | Hlnpv | -Eqs |
| 307 | 1.78 | 1.84 | 2.75 | 1.05 | 0.00 | 6 | Lkehm | -Qdr |
| 305 | 1.95 | 1.84 | 2.86 | 1.05 | 0.00 | 6 | Cdeinr | -Wgl |
| 311 | 1.78 | 2.24 | 2.90 | 1.05 | 0.00 | 6 | Trein | GSkpv |
| 533 | 0.82 | 0.00 | 1.47 | 1.05 | 0.00 | 5 | Kgq | - |
| 529 | 1.21 | 0.59 | 1.93 | 1.05 | 0.00 | 5 | Ifsv | -l |
| 654 | 0.81 | 1.38 | 1.97 | 1.05 | 0.00 | 5 | Sd | Ptn |
| 642 | 0.41 | 2.24 | 2.03 | 1.05 | 0.00 | 5 | Ed | I-lqs |
| 656 | 1.42 | 0.59 | 2.06 | 1.05 | 0.00 | 5 | Ivqt | Ph |
| 534 | 1.78 | 0.00 | 2.07 | 1.05 | 0.00 | 5 | Ivfkm | - |
| 532 | 1.21 | 1.15 | 2.14 | 1.05 | 0.00 | 5 | Aghs | -cf |
| 531 | 1.42 | 1.15 | 2.27 | 1.05 | 0.00 | 5 | Ivmn | -cq |
| 658 | 1.42 | 1.84 | 2.52 | 1.05 | 0.00 | 5 | Edkp | SAtv |
| 863 | 0.41 | 1.84 | 1.89 | 1.05 | 0.00 | 4 | Rt | QLav |
| 858 | 1.21 | 1.84 | 2.39 | 1.05 | 0.00 | 4 | Kamp | QEgl |
| 1116 | 0.65 | 0.00 | 1.36 | 1.05 | 0.00 | 3 | Kr | Q |
| 785 | 0.41 | 0.59 | 1.43 | 1.05 | 0.00 | 3 | Lv | -f |
| 872 | 0.41 | 0.99 | 1.57 | 1.05 | 0.00 | 3 | Ki | RQ |
| 1064 | 0.41 | 1.45 | 1.74 | 1.05 | 0.00 | 3 | Kt | QRe |
| 888 | 0.41 | 1.66 | 1.82 | 1.05 | 0.00 | 3 | Lv | Eadg |
| 890 | 0.41 | 2.13 | 1.99 | 1.05 | 0.00 | 3 | Sq | Gahkr |
| 1062 | 0.82 | 1.45 | 2.00 | 1.05 | 0.00 | 3 | Krv | DEl |
| 881 | 0.82 | 1.56 | 2.04 | 1.05 | 0.00 | 3 | Tav | -PS |
| 870 | 1.21 | 1.15 | 2.14 | 1.05 | 0.00 | 3 | Qeks | Rhn |
| 873 | 1.21 | 1.38 | 2.22 | 1.05 | 0.00 | 3 | Ndks | Eaq |
| 899 | 1.42 | 1.15 | 2.27 | 1.05 | 0.00 | 3 | Dept | -aq |
| 1144 | 1.42 | 1.15 | 2.27 | 1.05 | 0.00 | 3 | Lmft | Asv |
| 784 | 2.13 | 0.59 | 2.51 | 1.05 | 0.00 | 3 | Mldfiy | -n |
| 787 | 2.22 | 0.86 | 2.67 | 1.05 | 0.00 | 3 | RHdeks | -q |

| | | | | | | | | |
|---|---|---|---|---|---|---|---|---|
| 1143 | 1.78 | 1.66 | 2.69 | 1.05 | 0.00 | 3 | Aslnt | Rehq |
| 898 | 2.13 | 1.15 | 2.72 | 1.05 | 0.00 | 3 | Ysdkpr | -gl |
| 525 | 0.41 | 0.00 | 1.21 | 1.05 | 0.00 | 2 | Sm | - |
| 926 | 0.41 | 0.00 | 1.21 | 1.05 | 0.00 | 2 | Nt | K |
| 928 | 0.00 | 1.15 | 1.37 | 1.05 | 0.00 | 2 | V | Qae |
| 979 | 0.41 | 0.86 | 1.53 | 1.05 | 0.00 | 2 | Kl | Rq |
| 1107 | 1.04 | 0.00 | 1.61 | 1.05 | 0.00 | 2 | Edk | R |
| 1082 | 0.41 | 1.15 | 1.63 | 1.05 | 0.00 | 2 | Ed | Rqt |
| 1108 | 0.41 | 1.15 | 1.63 | 1.05 | 0.00 | 2 | Kr | Gns |
| 744 | 0.81 | 0.59 | 1.68 | 1.05 | 0.00 | 2 | Qn | -a |
| 1085 | 0.82 | 0.59 | 1.68 | 1.05 | 0.00 | 2 | Hay | Rq |
| 673 | 0.82 | 0.86 | 1.78 | 1.05 | 0.00 | 2 | Sct | Gr |
| 809 | 0.82 | 1.15 | 1.89 | 1.05 | 0.00 | 2 | Lfm | Ekp |
| 1086 | 0.41 | 1.84 | 1.89 | 1.05 | 0.00 | 2 | Qi | ASrv |
| 838 | 0.41 | 1.84 | 1.89 | 1.05 | 0.00 | 2 | Sd | PAgv |
| 793 | 0.41 | 1.84 | 1.89 | 1.05 | 0.00 | 2 | Rk | LTms |
| 526 | 1.58 | 0.00 | 1.95 | 1.05 | 0.00 | 2 | Slnpr | - |
| 683 | 0.81 | 1.38 | 1.97 | 1.05 | 0.00 | 2 | Dn | Rqe |
| 742 | 0.82 | 1.38 | 1.97 | 1.05 | 0.00 | 2 | Lfr | -pd |
| 980 | 1.42 | 0.59 | 2.06 | 1.05 | 0.00 | 2 | Stac | Wd |
| 681 | 0.82 | 1.66 | 2.08 | 1.05 | 0.00 | 2 | Qde | Flrv |
| 794 | 1.04 | 1.84 | 2.29 | 1.05 | 0.00 | 2 | Krh | QEns |
| 265 | 1.19 | 1.66 | 2.31 | 1.05 | 0.00 | 2 | V-k | Alrt |
| 1045 | 1.90 | 0.59 | 2.37 | 1.05 | 0.00 | 2 | Snah- | Dr |
| 1080 | 1.28 | 1.84 | 2.44 | 1.05 | 0.00 | 2 | SNl | KDat |
| 1016 | 2.13 | 0.59 | 2.51 | 1.05 | 0.00 | 2 | Saknt- | Di |
| 264 | 1.42 | 1.84 | 2.52 | 1.05 | 0.00 | 2 | R-ds | HQct |
| 820 | 2.52 | 1.38 | 3.05 | 1.05 | 0.00 | 2 | TAgpqsv | -ne |
| 1018 | 2.45 | 2.24 | 3.32 | 1.05 | 0.00 | 2 | Asfmpv- | EQgnt |
| 923 | 0.00 | 1.15 | 1.37 | 1.05 | 0.00 | 1 | D | Rhk |
| 1096 | 0.00 | 1.38 | 1.46 | 1.05 | 0.00 | 1 | K | Cfs |
| 709 | 0.00 | 1.38 | 1.46 | 1.05 | 0.00 | 1 | N | Edr |
| 1139 | 0.00 | 1.84 | 1.63 | 1.05 | 0.00 | 1 | K | AEst |
| 374 | 0.41 | 1.38 | 1.72 | 1.05 | 0.00 | 1 | E- | Rqh |
| 706 | 0.41 | 1.38 | 1.72 | 1.05 | 0.00 | 1 | Yf | Rqh |
| 702 | 0.00 | 2.13 | 1.73 | 1.05 | 0.00 | 1 | L | Qekrs |
| 687 | 0.98 | 0.59 | 1.79 | 1.05 | 0.00 | 1 | IL | Ag |
| 677 | 0.82 | 0.99 | 1.83 | 1.05 | 0.00 | 1 | Hen | KR |
| 948 | 0.82 | 1.15 | 1.89 | 1.05 | 0.00 | 1 | Yn- | Qcr |
| 406 | 0.41 | 1.95 | 1.93 | 1.05 | 0.00 | 1 | Ne | AGVm |
| 724 | 0.82 | 1.38 | 1.97 | 1.05 | 0.00 | 1 | Ils | Evg |
| 945 | 0.82 | 1.38 | 1.97 | 1.05 | 0.00 | 1 | Acy | Ltg |
| 664 | 0.82 | 1.45 | 2.00 | 1.05 | 0.00 | 1 | Fay | LMq |
| 670 | 0.82 | 1.66 | 2.08 | 1.05 | 0.00 | 1 | Rms | Qchk |
| 1148 | 0.82 | 1.84 | 2.14 | 1.05 | 0.00 | 1 | Edl | PRgs |
| 974 | 1.25 | 1.15 | 2.16 | 1.05 | 0.00 | 1 | Hkq | Liv |
| 908 | 0.82 | 2.24 | 2.29 | 1.05 | 0.00 | 1 | -dv | IMalt |
| 893 | 0.82 | 2.24 | 2.29 | 1.05 | 0.00 | 1 | Iem | LPgqt |
| 1058 | 1.42 | 1.45 | 2.38 | 1.05 | 0.00 | 1 | Rkes | ATq |
| 328 | 1.21 | 1.84 | 2.39 | 1.05 | 0.00 | 1 | Lcs- | GEpr |
| 397 | 1.78 | 0.86 | 2.39 | 1.05 | 0.00 | 1 | Mqikl | As |
| 633 | 1.58 | 1.38 | 2.46 | 1.05 | 0.00 | 1 | Sadin | R-p |
| 849 | 1.58 | 1.38 | 2.46 | 1.05 | 0.00 | 1 | Tlmnv | Kap |
| 812 | 1.21 | 2.13 | 2.50 | 1.05 | 0.00 | 1 | Scnv | Akpqt |
| 798 | 1.21 | 2.13 | 2.50 | 1.05 | 0.00 | 1 | Ndsy | Kaeqt |
| 435 | 1.58 | 1.66 | 2.56 | 1.05 | 0.00 | 1 | Qahkn | Lmsy |
| 471 | 1.78 | 1.38 | 2.58 | 1.05 | 0.00 | 1 | Lngrt | Asd |
| 62 | 1.78 | 1.38 | 2.58 | 1.05 | 0.00 | 1 | Smhl- | Gaw |
| 1151 | 1.58 | 1.84 | 2.63 | 1.05 | 0.00 | 1 | Rhint | -Plv |
| 1128 | 1.42 | 2.24 | 2.67 | 1.05 | 0.00 | 1 | Vtdy | ESaqr |
| 271 | 1.73 | 1.84 | 2.72 | 1.05 | 0.00 | 1 | Nseg | AKrt |
| 324 | 1.95 | 1.66 | 2.79 | 1.05 | 0.00 | 1 | Lfqst- | Parw |
| 474 | 1.83 | 2.24 | 2.93 | 1.05 | 0.00 | 1 | APTq | RSilm |
| 606 | 0.82 | 1.84 | 2.04 | 0.94 | 0.11 | 7 | Iln | D-lq |
| 306 | 1.58 | 2.13 | 2.63 | 0.94 | 0.11 | 6 | Denpy | -aekv |
| 309 | 1.95 | 2.24 | 2.90 | 0.94 | 0.11 | 6 | Idfgmt | A-npt |
| 660 | 0.82 | 1.66 | 1.97 | 0.94 | 0.11 | 5 | Kir | Eaiq |
| 644 | 1.21 | 1.38 | 2.11 | 0.94 | 0.11 | 5 | Flvy | G-l |
| 659 | 1.58 | 1.15 | 2.27 | 0.94 | 0.11 | 5 | Saknt | Pgt |
| 645 | 1.21 | 1.84 | 2.29 | 0.94 | 0.11 | 5 | Qhkn | S-ch |
| 860 | 0.41 | 2.13 | 1.89 | 0.94 | 0.11 | 4 | De | Vaekl |
| 862 | 1.58 | 2.24 | 2.67 | 0.94 | 0.11 | 4 | Qaklr | FGatv |
| 1142 | 0.82 | 1.15 | 1.78 | 0.94 | 0.11 | 3 | Itv | Alv |

| | | | | | | | | |
|---|---|---|---|---|---|---|---|---|
| 889 | 0.41 | 2.13 | 1.89 | 0.94 | 0.11 | 3 | Ed | Padls |
| 1024 | 1.42 | 1.15 | 2.16 | 0.94 | 0.11 | 3 | Edt- | Gk- |
| 880 | 1.42 | 1.15 | 2.16 | 0.94 | 0.11 | 3 | Lmrv | Igv |
| 1114 | 1.21 | 1.66 | 2.22 | 0.94 | 0.11 | 3 | Qgkr | Eadk |
| 1115 | 1.78 | 1.15 | 2.39 | 0.94 | 0.11 | 3 | Ntals | Eaq |
| 882 | 1.55 | 1.84 | 2.50 | 0.94 | 0.11 | 3 | Mvip | -Qlp |
| 1021 | 1.42 | 2.13 | 2.52 | 0.94 | 0.11 | 3 | Sta- | Wghm- |
| 1023 | 1.95 | 1.45 | 2.61 | 0.94 | 0.11 | 3 | Acpst- | GQ- |
| 36 | 1.04 | 1.38 | 2.01 | 0.94 | 0.11 | 2 | G-s | Aps |
| 912 | 0.82 | 2.13 | 2.14 | 0.94 | 0.11 | 2 | Lfv | Iamnv |
| 911 | 1.42 | 1.15 | 2.16 | 0.94 | 0.11 | 2 | Rkde | Adn |
| 819 | 1.42 | 1.84 | 2.42 | 0.94 | 0.11 | 2 | Vsmp | -Qpr |
| 963 | 1.90 | 1.15 | 2.47 | 0.94 | 0.11 | 2 | Tsgin | Agv |
| 807 | 2.29 | 1.66 | 2.91 | 0.94 | 0.11 | 2 | Mhklnqt | Aksv |
| 1099 | 0.41 | 0.59 | 1.32 | 0.94 | 0.11 | 1 | Ny | Hy |
| 1136 | 0.82 | 1.38 | 1.87 | 0.94 | 0.11 | 1 | Rdq | Aeq |
| 628 | 0.82 | 1.66 | 1.97 | 0.94 | 0.11 | 1 | Eks | Hdps |
| 277 | 1.21 | 1.15 | 2.03 | 0.94 | 0.11 | 1 | Kadg | Rgh |
| 639 | 0.82 | 1.84 | 2.04 | 0.94 | 0.11 | 1 | Nht | S-gh |
| 969 | 0.82 | 1.84 | 2.04 | 0.94 | 0.11 | 1 | Qsy | PAst |
| 1092 | 0.82 | 1.84 | 2.04 | 0.94 | 0.11 | 1 | Vim | TNis |
| 1104 | 1.58 | 0.59 | 2.06 | 0.94 | 0.11 | 1 | Aeins | Ls |
| 790 | 1.42 | 1.38 | 2.25 | 0.94 | 0.11 | 1 | Dhen | -ae |
| 1072 | 1.21 | 1.84 | 2.29 | 0.94 | 0.11 | 1 | Sefk | RAel |
| 44 | 1.95 | 0.59 | 2.29 | 0.94 | 0.11 | 1 | Denqs- | G- |
| 366 | 1.55 | 1.38 | 2.33 | 0.94 | 0.11 | 1 | Ken- | Rq- |
| 466 | 1.42 | 1.66 | 2.35 | 0.94 | 0.11 | 1 | Spdq | Ldtv |
| 905 | 1.78 | 1.15 | 2.39 | 0.94 | 0.11 | 1 | Gpils | Ael |
| 254 | 1.78 | 1.45 | 2.50 | 0.94 | 0.11 | 1 | Nsdi- | GH- |
| 240 | 1.58 | 1.84 | 2.52 | 0.94 | 0.11 | 1 | Ndel- | GAh- |
| 695 | 1.58 | 1.84 | 2.52 | 0.94 | 0.11 | 1 | Nikqy | REdy |
| 601 | 1.90 | 1.38 | 2.55 | 0.94 | 0.11 | 1 | Vifnt | Plt |
| 41 | 2.13 | 1.15 | 2.61 | 0.94 | 0.11 | 1 | Y-ahkn | Sat |
| 816 | 1.95 | 1.66 | 2.69 | 0.94 | 0.11 | 1 | Adhks- | Geis |
| 339 | 1.95 | 1.84 | 2.75 | 0.94 | 0.11 | 1 | Siklv- | GPir |
| 148 | 1.78 | 2.13 | 2.75 | 0.94 | 0.11 | 1 | Qrky- | Ehps- |
| 260 | 1.58 | 2.52 | 2.77 | 0.94 | 0.11 | 1 | Sakr- | Qaelmn |
| 200 | 1.78 | 2.52 | 2.90 | 0.94 | 0.11 | 1 | Yfce- | Amqvw- |
| 318 | 2.13 | 2.24 | 3.01 | 0.94 | 0.11 | 1 | V-afkm | DTals |
| 648 | 0.41 | 1.38 | 1.57 | 0.90 | 0.14 | 5 | L- | P-s |
| 646 | 1.21 | 1.38 | 2.08 | 0.90 | 0.14 | 5 | Tag- | L-s |
| 109 | 0.82 | 1.38 | 1.83 | 0.90 | 0.14 | 3 | Ht- | D-r |
| 111 | 0.82 | 1.38 | 1.83 | 0.90 | 0.14 | 3 | Aq- | V-p |
| 110 | 1.42 | 0.86 | 2.02 | 0.90 | 0.14 | 3 | Lsi- | P- |
| 1061 | 1.04 | 2.24 | 2.29 | 0.90 | 0.14 | 3 | Knq | LQeir |
| 896 | 1.58 | 2.24 | 2.63 | 0.90 | 0.14 | 3 | Limqt | M-gkr |
| 840 | 1.04 | 1.56 | 2.04 | 0.90 | 0.14 | 2 | Tls | PAS |
| 964 | 1.21 | 1.56 | 2.14 | 0.90 | 0.14 | 2 | Rkqy | LHQ |
| 674 | 1.19 | 1.84 | 2.23 | 0.90 | 0.14 | 2 | Rkc | SCnt |
| 34 | 1.78 | 1.84 | 2.61 | 0.90 | 0.14 | 2 | N-irs | ASel |
| 1046 | 1.90 | 1.84 | 2.68 | 0.90 | 0.14 | 2 | Edrs- | PSnt |
| 934 | 0.82 | 1.38 | 1.83 | 0.90 | 0.14 | 1 | Mtv | Avd |
| 114 | 1.58 | 1.38 | 2.31 | 0.90 | 0.14 | 1 | Adkt- | S-p |
| 738 | 1.21 | 2.24 | 2.39 | 0.90 | 0.14 | 1 | Qelm | LStv- |
| 1011 | 1.58 | 1.95 | 2.52 | 0.90 | 0.14 | 1 | Caipv | AEGs |
| 1027 | 1.95 | 1.38 | 2.54 | 0.90 | 0.14 | 1 | Teinp- | A-v |
| 1055 | 2.05 | 1.38 | 2.61 | 0.90 | 0.14 | 1 | NEdgs | Asr |
| 996 | 1.95 | 1.84 | 2.71 | 0.90 | 0.14 | 1 | Lakmsv | GSpr |

**Copyright (c) Anton Feenstra 2007**
